# A site-specific analysis of the ADP-Ribosylome unveils Homogeneous DNA Damage-Induced Serine ADP-Ribosylation across wild-type and BRCA-mutant Breast Cancer cell lines

**DOI:** 10.1101/2023.12.15.571635

**Authors:** Holda A. Anagho, Meeli Mullari, Aurél G. Prósz, Sara C. Buch-Larsen, Marie Locard-Paulet, Zoltan Szallasi, Michael L. Nielsen

## Abstract

ADP-ribosylation (ADPr) signaling plays a crucial role in the DNA damage response. Inhibitors against the main enzyme catalyzing ADPr after DNA damage – PARP1 – are used as targeted therapies against breast cancers with BRCA1/2 mutations. However, development of resistance to PARP inhibitors (PARPi) is a major obstacle in treating patients. To better understand the role of ADPr in PARPi sensitivity, we used Liquid Chromatography-Mass Spectrometry (LC-MS) for systems level analysis of the ADP-ribosylome in six breast cancer cell lines exhibiting different PARPi sensitivities. We identified 1,632 sites on 777 proteins across all cell lines, primarily on serine residues, with site-specific overlap of targeted residues across DNA damage-related proteins across all cell lines, demonstrating high conservation of serine ADPr signaling networks upon DNA damage. We furthermore observed site-specific differences in ADPr intensities in PARPi-sensitive BRCA mutants, and unique ADPr sites in PARPi-resistant BRCA mutant cells, which we notably show to have low PARG levels and longer ADPr chains on PARP1.

## Introduction

Protein post-translational modifications (PTMs) act as molecular switches modulating various cellular processes and add significant complexity to the proteome^1^. Perturbation of the enzymes catalyzing PTM addition or removal can therefore disrupt signaling networks and lead to diseases, making these enzymes important targets for therapeutic interventions^2^. Systems-level investigation of PTMs provides insights to understand regulatory mechanisms behind protein changes in human cells^3-7^. This approach also helps identify PTMs that play crucial roles in cellular processes, as these tend to be conserved across different cell types^8-10^.

ADP-ribosylation (ADPr) is the post-translational addition of adenosine dinucleotide phosphate-ribose onto amino acid residues on target proteins. The reaction is carried out by a diverse group of enzymes collectively referred to as ADP-ribosyltransferases (ARTs) or Poly (ADP-ribose) polymerase (PARP), using the redox molecule nicotinamide adenine dinucleotide (NAD^+^) as a substrate. Proteins can be mono-ADP-ribosylated (MARylated) or poly-ADP-ribosylated (PARylated) in linear or branched chains on a variety of residues including Arginine (R), Aspartic acid (E), Cysteine (C), Glutamic acid (D), Histidine (H), Lysine (K), Serine (S), Threonine (T), or Tyrosine (Y)^11^. ADPr groups are removed from target proteins by glycohydrolases, including poly ADP-ribose glycohydrolase (PARG), which hydrolyzes the ribose-ribose bonds within poly-ADPr (PAR) chains, and ADP-ribosylhydrolase 3 (ARH3), which hydrolyzes serine-ADPr bonds^12,13^. Because ADP-ribose is a bulky, negatively charged molecule, ADPr of a protein can modulate its activity or interactions with DNA or other proteins^14^.

Consequently, ADPr is implicated in most major cellular processes, including the DNA damage response^15^. In this context, ADPr is primarily mediated by PARP1 with a cofactor HPF1^16,17^. Upon DNA damage, PARP1 can bind single strand breaks (SSBs) and double-strand breaks (DSBs), which induces ADPr of itself and downstream target proteins involved in repairing the damaged DNA^18-22^. Two other proteins essential for DNA damage repair are BRCA1 and BRCA2, with studies published in 2005 reporting that loss of BRCA1 or BRCA2 function sensitized cells to PARP inhibition, suggesting a synthetic lethal interplay between the two genes^23,24^. As a result, PARP inhibitors (PARPi) Olaparib, Rucaparib, and Talazoparib are now used in the clinic for the treatment of breast, ovarian, and prostate cancers with germline BRCA mutations, and clinical trials for patients with other HRR defects is ongoing^25-28^. Notwithstanding their clinical efficacy, the emergence of resistance to PARP inhibitors (PARPi) poses a significant challenge in the long-term treatment of cancer patients^29^. Hence, understanding how ADPr is regulated across breast cancer may provide insights into the mechanisms leading to PARPi resistance and further the development of novel treatment strategies.

Although the importance of PARP1 in DNA damage repair (DDR) signaling is established, the exact mechanistic details are less well understood. Mass spectrometry (MS)-based characterization of ADPr sites has emerged as a valuable methodology towards elucidating the molecular details related to the functional role of PARP1 and ADPr in DDR^30^. Recently, a study reported a proteome-wide characterization of acid-linked (D/E) ADPr in a panel of breast cancer cell lines^31^. Here, the authors reported substantial heterogeneity in D/E-ADPr across the cell lines, attributed to cell line-specific PARP1 activation in breast cancer. However, this study did not account for serine ADPr, which has recently been shown to be a major target of PARP1 in DNA damage^16,32-35^.

Therefore, to get a more complete picture of DNA damage-induced ADPr, we used MS to conduct a systems-level profiling of ADPr in breast cancer cell lines. Collectively, we identified 1,632 ADPr sites on 777 proteins, with 92% of ADPr events occurring on serine residues. We found that H_2_O_2_ treatment induced robust and homogeneous ADPr of DNA damage repair proteins in all cell lines, regardless of BRCA1/2 mutation or PARPi sensitivity. Furthermore, ADPr sites and proteins were conserved across different cell lines^35^ and organisms^36^. Interestingly, we found reduced PARG expression in PARPi resistant BRCA1 mutant cells, and we found that ADPr on USF1 was significantly upregulated in PARPi sensitive cells compared to PARPi resistant cells. These findings provide important evidence for the existence of a homogeneous ADPr signaling network centered on serine ADPr as a biologically significant PTM. Collectively, our findings will serve as a useful resource for investigating the roles of site-specific serine ADPr in DNA damage repair, which hitherto has been remained unexplored.

## Results

### Breast cancer cells express different levels of ADPr signaling enzymes and have different PARP inhibitor sensitivities

For this study, we chose six cell lines representing a variety of breast cancer subtypes, tumor origins, and molecular markers (**Table 1**), including three BRCA1/2 wild type, two BRCA1 mutants, and one BRCA2 mutant. We first used two whole-genome sequencing-based classifiers, Homologous Recombination Deficiency (HRD) Score and HRDetect^37,38^ to predict Homologous recombination repair (HRR) deficiency in the cell lines and compared them to 45 other breast cancer cell lines based on whole exome sequences from the Cancer Cell Line Encyclopedia (CCLE). HRD score was developed to identify triple-negative breast cancer (TNBC) tumors, including BRCA1/2 wild-type tumors, likely to respond to DNA-damaging platinum agents, whereas HRDetect was trained on breast cancer patient samples to predict BRCA1/2 deficiency as a proxy for PARPi sensitivity.

As expected, the BRCA mutant cell lines MDAMB436, HCC1937 and HCC1428 scored above the cutoff for HRD Score (>42)^39^ and HRDetect (>0.7)^40^, supporting their functional BRCA deficiency and thereby HRR defects (**Fig. 1A**). Conversely, BRCA WT cell lines T47D and MDAMB231 scored below the cutoff for HRDetect, while T47D was just at the HRD score cutoff. Surprisingly, MCF7 scored well above the cutoff values for both HRD Score and HRDetect score, suggesting HRR deficiency. However, this observation could possibly be explained by a chromosomal rearrangement in this cell line resulted in an in-frame fusion of the *RAD51C* gene with the *ATXN7* gene^41^. Based on these results, MDAMB436, HCC1937, and HCC1428 were predicted to be more sensitive to PARP inhibitors compared to MDAMB231, T47D, and MCF7, in line with published reports^42^.

**Figure 1.**
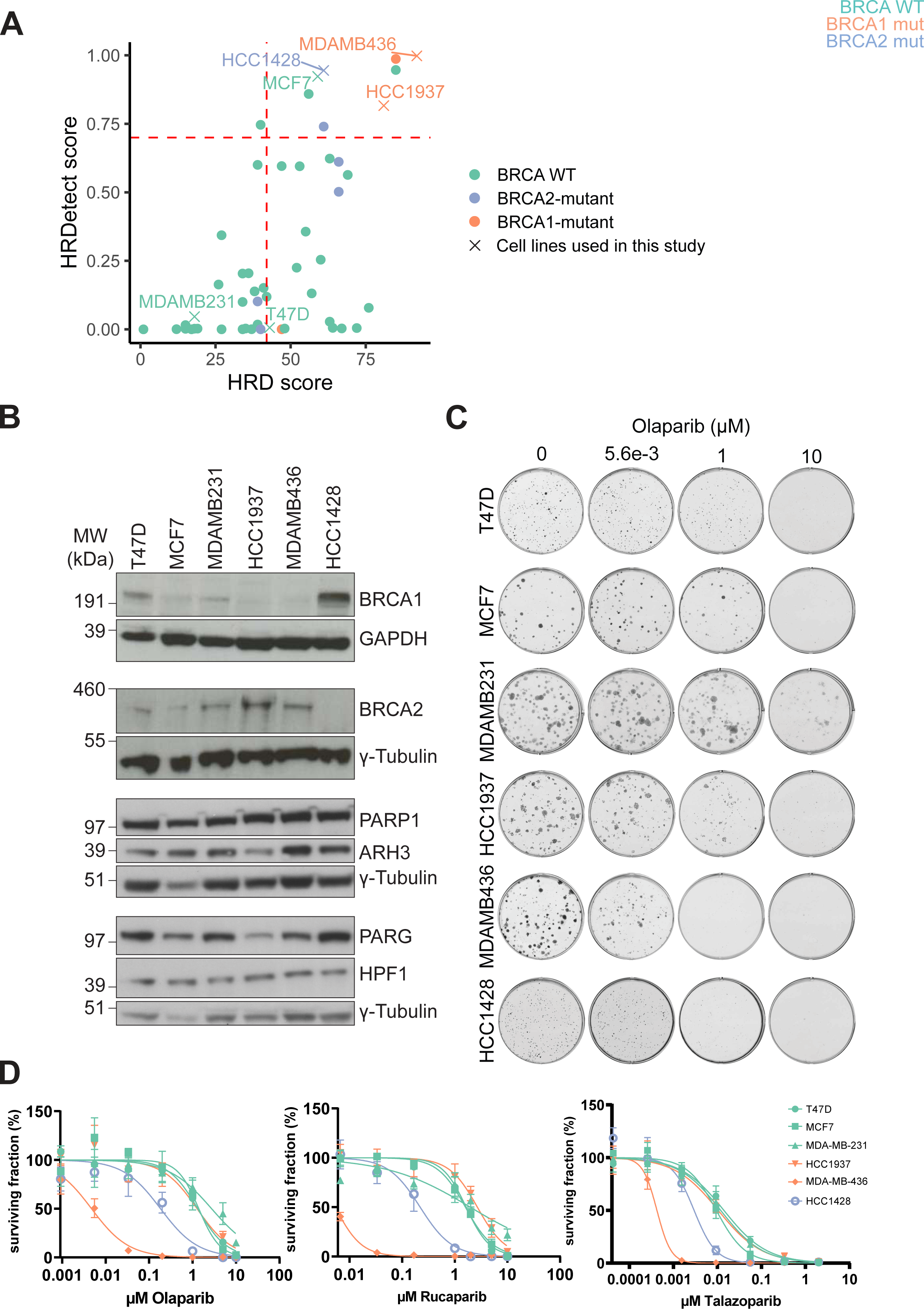
Different sensitivity to PARPi and expression of ADPr signaling enzymes among BRCA mutant cell lines. **A.** The logistic regression-based classifiers HRD and HRDetect score predicting homologous recombination repair deficiency are shown for 51 breast cancer cell lines. The dotted lines indicate the threshold for predicted homologous recombination deficiency. **B.** Western blots show expression of BRCA1 and BRCA2, ADPr writer enzyme PARP1 and its cofactor HPF1, and eraser enzymes PARG and ARH3 in cell lines. **C-D**. Colony formation assays were performed on breast cancer cell lines treated with a range of concentrations of indicated PARPi for 9 – 18 days. Colony formation in Olaparib-treated cells is shown in **C**, and percent survival relative to DMSO controls is shown in **D**. Error bars represent the standard deviation of 3 replicate wells per drug concentration. Average CC_50_ values for Olaparib, Rucaparib, and Talazoparib are shown in Table 2. Cytotoxicity curves and average CC_50_ values were calculated in Graphpad Prism, using the [Inhibitor] vs. normalized response – Variable slope nonlinear fit formula.

To further this, we next tested the sensitivity of the cell lines to the clinically approved PARPi Olaparib, Rucaparib, or Talazoparib using colony formation assays (**Fig. 1C&1D**) and calculated the amount of each PARPi required to inhibit the growth of 50% of the colonies CC_50_ **(Table 2**). From this, we found that BRCA1 mutant MDAMB436 cells were the most sensitive to all three PARPi, followed by BRCA2 mutant HCC1428 cells, in agreement with our HRD Score and HRDetect Score predictions. Interestingly, BRCA1 mutant HCC1937 cells were not sensitive to Olaparib, Rucaparib, or Talazoparib compared to the BRCA wild-type cell lines, contrary to our HRD score and HRDetect score predictions, but in line with previous observations^43^. Furthermore, we did not observe PARPi sensitivity in MCF7 cells compared to the other BRCA wild type cells.

Next, we investigated the expression levels of BRCA1 and BRCA2 in the relevant cell lines by western blot (WB), with MCF7 cells having the lowest levels compared to the other BRCA1-expressing cell lines (**Fig. 1B**). To explore whether the cell lines express the enzymes relevant for DNA damage-induced serine ADPr signaling stimulated by H_2_O_2_ treatment, we analyzed expression levels of the major ADPr writer enzyme PARP1, its cofactor HPF1, and main erasers PARG and ARH3 by WB (**Fig. 1B**). We confirmed that all cell lines express the proteins needed for serine ADPr signaling after DNA damage with lowest expression of ARH3 and PARG in HCC1937 cells, and highest expression of ARH3 in MDAMB436 cells.

### MS-based proteome profiling of breast cancer cell lines shows differential expression of ADPr signaling enzymes in BRCA mutant cell lines with different PARPi sensitivity

Having established the individual cell lines response to PARPi, we next wanted to explore the overall protein expression differences across the cell lines by characterizing their individual proteomes. To this end, we used a standard reverse-phase high pH liquid chromatography fractionation strategy^44^ with each fraction subsequently analyzed using data-dependent acquisition (DDA) shotgun proteomics workflow **(Fig. 2A**). All experiments were performed in quadruplicate with quantification of individual proteins obtained using label-free quantification (LFQ)^45^. Collectively, we identified 226,624 peptides in total and quantified around 7000 unique proteins in each cell line (**Supp. Fig. 1A-B; Supp. Table 1**). LFQ intensities across all samples were similar, CVs were below 1% between replicates, and the Pearson correlations between replicates were (>0.96) (**Supp. Fig. 1D-F**), demonstrating high reproducibility between replicates.

**Figure 2.**
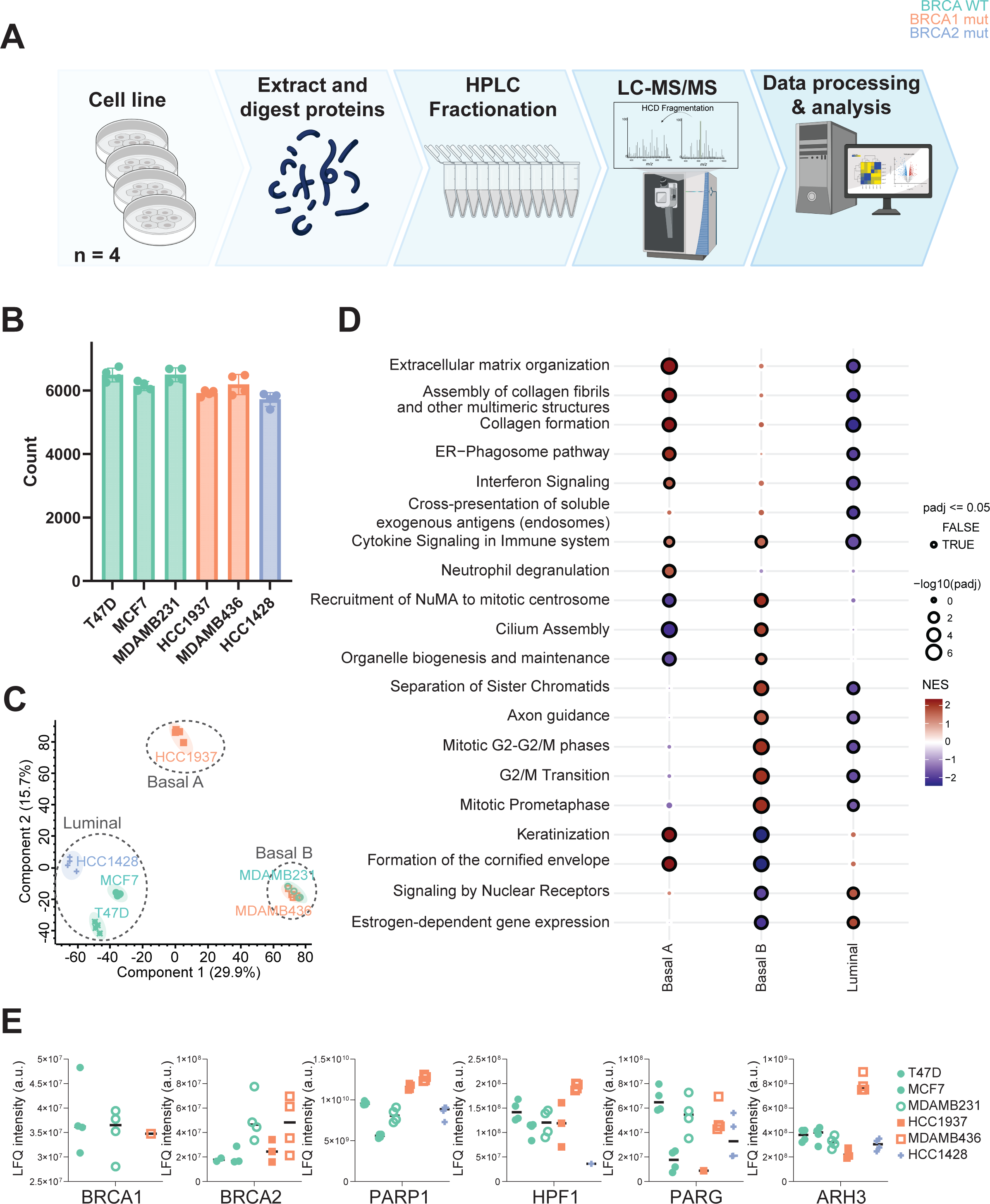
The proteomes of breast cancer cell lines shows cluster by breast cancer molecular subtype and confirms differential expression of ADPr signaling enzymes among BRCA mutant cell lines. **A.** Overview of the workflow used for proteome profiling of breast cancer cell lines. Cells were lysed in Guanidine hydrochloride (GndHCl) buffer, digested with Lys-C and Trypsin, separated into 12 concatenated fractions using high pH liquid chromatography, and analyzed by LC-MS/MS. Each cell line was analyzed in quadruplicate. **B** Number of proteins identified in each replicate of each cell line. **C.** Principal component analysis reveals clustering of cell lines according to molecular subtype. **D.** Differential expression analysis followed by gene set enrichment analysis (GSEA) with Reactome terms was performed on the proteins identified in Basal A (HCC1937), Basal B (MDAMB231 and MDAMB436) and Luminal (T47D, MCF7, HCC1428) cells using fGSEA^47^, and the top 20 differentially regulated Reactome pathways in each subtype relative to the other subtypes were plotted on a dotplot. The size of each circle represents the -log10 of the adjusted p-Value (padj), and statistical significance (padj <=0.05) is indicated with a black circle. Dots are colored by Normalized Enrichment Score (NES), with positive scores in red and negative scores in blue. **E.** Expression of BRCA1, BRCA2, PARP1, HPF1, PARG, and ARH3 in cell lines was detected by mass spectrometry.

As expected, principal component analysis (PCA) showed that the cell lines cluster according to their luminal (HCC1428, T47D, MCF7), basal A (HCC1937), and basal B (MDAMB231, MDAMB436) molecular subtypes (**Fig. 2D**) determined from microarray analyses of cell line transcriptomes^46^. Gene Set Enrichment Analysis (GSEA), where LFQ intensities for each protein in a sample group is compared to all other sample groups, was used to elucidate pathways significantly up- or down-regulated in the different cell types and in the different molecular subtypes^47^. We found that estrogen-dependent gene expression was upregulated in Luminal cells, consistent with the reported Estrogen Receptor-(ER) and Progesterone Receptor-(PR) positive nature of these cell lines^46^, while it was downregulated in ER- and PR-negative Basal B cells. Furthermore, pathways related to extracellular matrix formation and expression of structural and fibrous proteins were significantly upregulated in Basal A cells and downregulated in Basal B and luminal cells, while mitotic pathways are significantly upregulated in Basal B cells and downregulated in Luminal cells. Comparison between individual cell lines showed that a diverse range of pathways were significantly upregulated or downregulated in each cell line compared to the other five (**Supp. Fig. 1F**). For example, mitochondrial translation pathways are significantly downregulated in HCC1428 cells, whereas mitosis pathways were upregulated. On the other hand, mitochondrial translation pathways are down in MDAMB231 cells, whereas pathways related to mitosis are upregulated.

The ADPr signaling enzymes PARP1, HPF1, PARG, and ARH3 were quantified by MS across the samples, and PARP1 expression levels were similar across cell lines. On the other hand, PARG and ARH3 protein levels exhibited lowest expression in the PARPi resistant BRCA1 mutant cell line, while HPF1 protein expression levels were lowest in the HCC1428 cell line, whereas all three enzymes had very high expression in the most PARPi sensitive MDAMB436 cell line (**Fig. 2E**). These results support that although the proteomes of the breast cancer cell lines cluster according to molecular subtypes, there are differences in the expression of ADPr signaling proteins in the BRCA mutant cell lines exhibiting different PARPi sensitivity.

### ADP-ribosylation in breast cancer cells is strongly induced by H_2_O_2_ treatment and is predominantly serine-targeted

For site-specific characterization of ADPr in the breast cancer cell lines, we first treated the cells with 1 mM H_2_O_2_ for ten minutes to induce DNA damage and PARP1 activation^48^. The cells were then subjected to Af1521-based ADPr enrichment and EThcD mass spectrometry as previously described (Fig. 3A)^35,49,50^. Strong induction of ADPr by H_2_O_2_ treatment was confirmed both by MS and WB analysis in all six breast cancer cell lines (**Fig. 3B-C, Supp. Fig. 2A**). The median CVs in replicates from each cell line remained below 10% (**Supp. Fig. 2B**).

**Figure 3.**
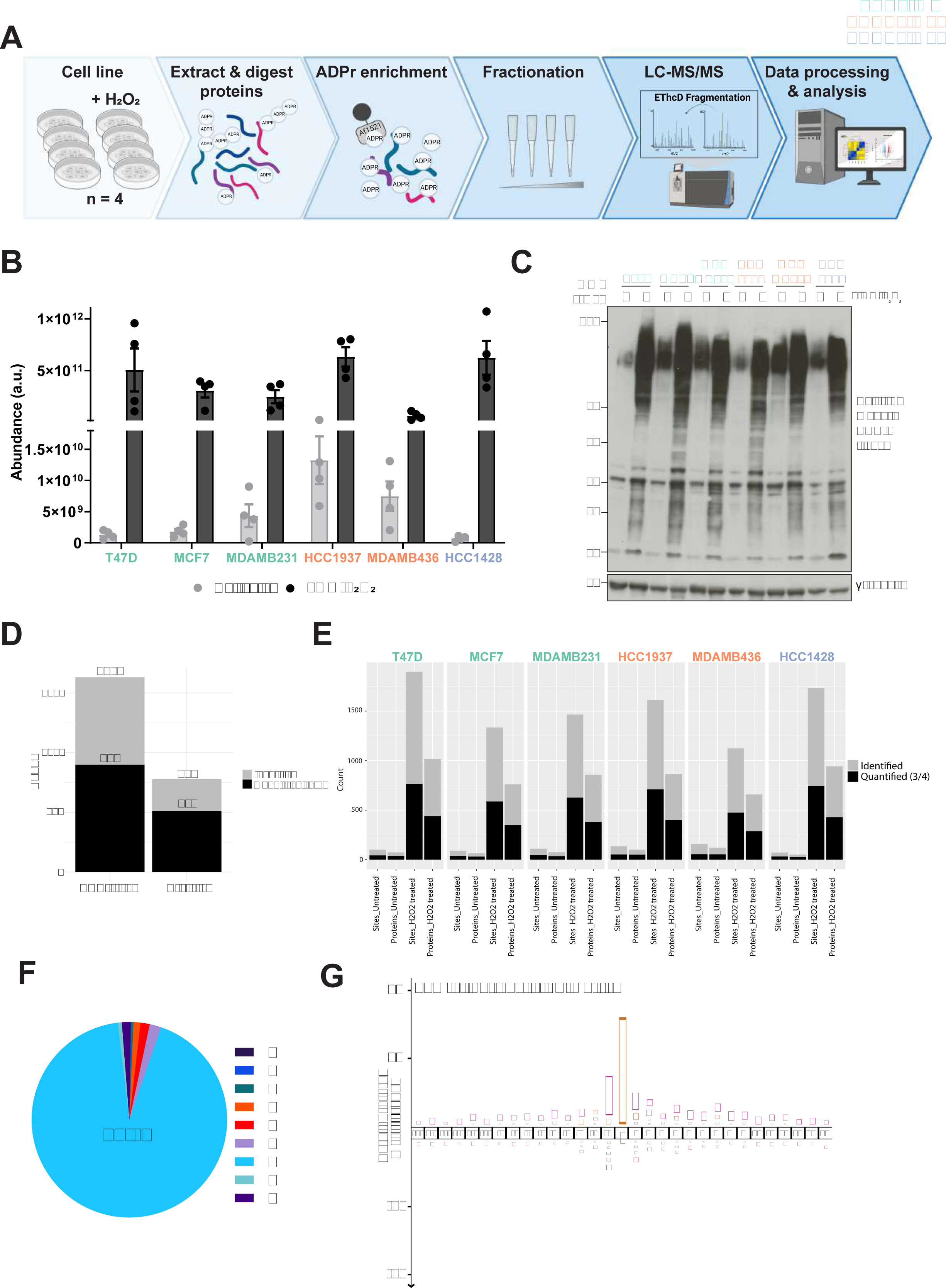
Serine ADP-ribosylation is strongly stimulated in breast cancer cell lines in response to H_2_O_2_ treatment. **A**. Overview of the workflow used for mass spectrometry characterization of ADPr in breast cancer cell lines. Cells were treated with 1 mM H_2_O_2_ for 10 minutes, lysed in GndCl and digested into peptides, purified and lyophilized to facilitate concentration. Peptides were incubated with PARG to digest poly-ADP-ribose chains into mono-ADPr moieties, and ADP-ribosylated peptides were enriched with Af1521 macrodomains. Peptides were cleaned up and fractionated on stage tips and analyzed by LC-MS/MS with electron-transfer/higher-energy collusion dissociation (EThcD). Each sample was prepared in quadruplicate. **B-E.** Induction of ADP-ribosylation in all cell lines after H_2_O_2_ treatment. Abundance of ADPr peptides before and after H_2_O_2_ treatment identified by mass spectrometry **(B)** and ADPr proteins by western blot **(C). D.** The bar chart shows the total number of ADP-ribosylated sites and proteins identified in total with a localization score >0.90 (gray) or quantified in 3 or more replicates of at least one sample (black). **E.** The bar chart shows the total number of ADP-ribosylated sites and proteins in untreated or H_2_O_2_-treated breast cancer cells. Sites or proteins identified with a localization score >0.90 are shown in gray while those quantified in 3 or more replicates of at least one sample are in black. **F.** Pie chart of the amino acid distribution of ADPr target sites shows that Serine is the major ADPr target residue. **G.** IceLogo generated from sequence windows surrounding the 1515 serine-ADPr sites identified, with the human Swiss-Prot proteome as a reference set. Amino acids above the line are enriched whereas amino acids below the line are de-enriched. Workflow figure was made with biorender.com.

In total, we identified 1,632 unique ADPr sites on 777 proteins with localization probability >0.9 (**Fig. 3D, Supp. Table 2**). In untreated cells, we identified the most ADPr events in MDAMB436 cells (106 sites on 67 proteins), followed by HCC1937 cells (86 sites on 51 proteins) (**Fig 3E**). By contrast, among the H_2_O_2_-treated cells, the largest number of ADPr sites were identified in T47D cells (1,133 sites on 576 proteins) and the lowest in MDAMB436 cells (649 sites on 371 proteins). As previously described in HeLa cells as well as *Drosophila* cell lines^35,36^, serine emerged as a major ADPr target amino acid, with 92.8% of the total identified ADPr acceptor sites localizing to serine residues (**Fig. 3F**). In agreement with previous reports^33^, we observed a strong KS motif adherence in 53% of serine-ADPr sites (**Fig. 3G**). Before H_2_O_2_ treatment, all ADPr samples cluster somewhat together in a PCA plot, while the HCC1937 and MDAMB436 separated slightly from the other samples **(Supp. Fig. 2C**). Interestingly, after H_2_O_2_ treatment, the HCC1937 ADPr samples cluster separately from all the other cell lines, suggesting a difference in the ADPr response in response to DNA-damage for this PARPi resistant BRCA mutant cell line. (**Supp. Fig. 2D**). Together, these results show that H_2_O_2_ strongly stimulates ADPr in all breast cancer cell lines, with a strong preference for serine residues.

### DNA damage triggers a homogenous serine ADP-ribosylation response across PARPi sensitive and resistant breast cancer cells

In total, we were able to quantify 899 ADPr sites in the H_2_O_2_-treated breast cancer cell lines on 511 proteins (**Fig. 3D**), which excludes low abundant sites that were detected in fewer than 3 replicates (**Supp. Fig. 3A&3B**). Among these, 317 ADPr sites were present in all six cell lines, and an additional 173 sites identified in five out of the six cell lines, suggesting that identified serine ADPr modification sites are homogenously targeted across different breast cancer cell lines, in contrast to previous observations related to D/E-ADPr^31^ (**Fig. 4A and Supp. Fig. 3C**). Consistent with the site level overlap, there was also a high ADPr protein level overlap between the cell lines (**Supp. Fig. 3D**). Performing GSEA using Enrichr^51^ on the ADPr target proteins containing the 317 ADPr sites quantified in all six cell lines showed an enrichment of proteins involved in DNA repair, DNA damage/Telomere Stress induced senescence, Base Excision Repair, and DNA double-strand break repair, and Epigenetic regulation of gene expression (**Fig. 4B**). Network analysis and functional enrichment of the ADPr target proteins found in all six cell lines^52^ further showed that these proteins are involved in DNA repair, RNA metabolism or Ribosome Biogenesis, or Nucleosome components including histones (**Fig. 4C**).

**Figure 4:**
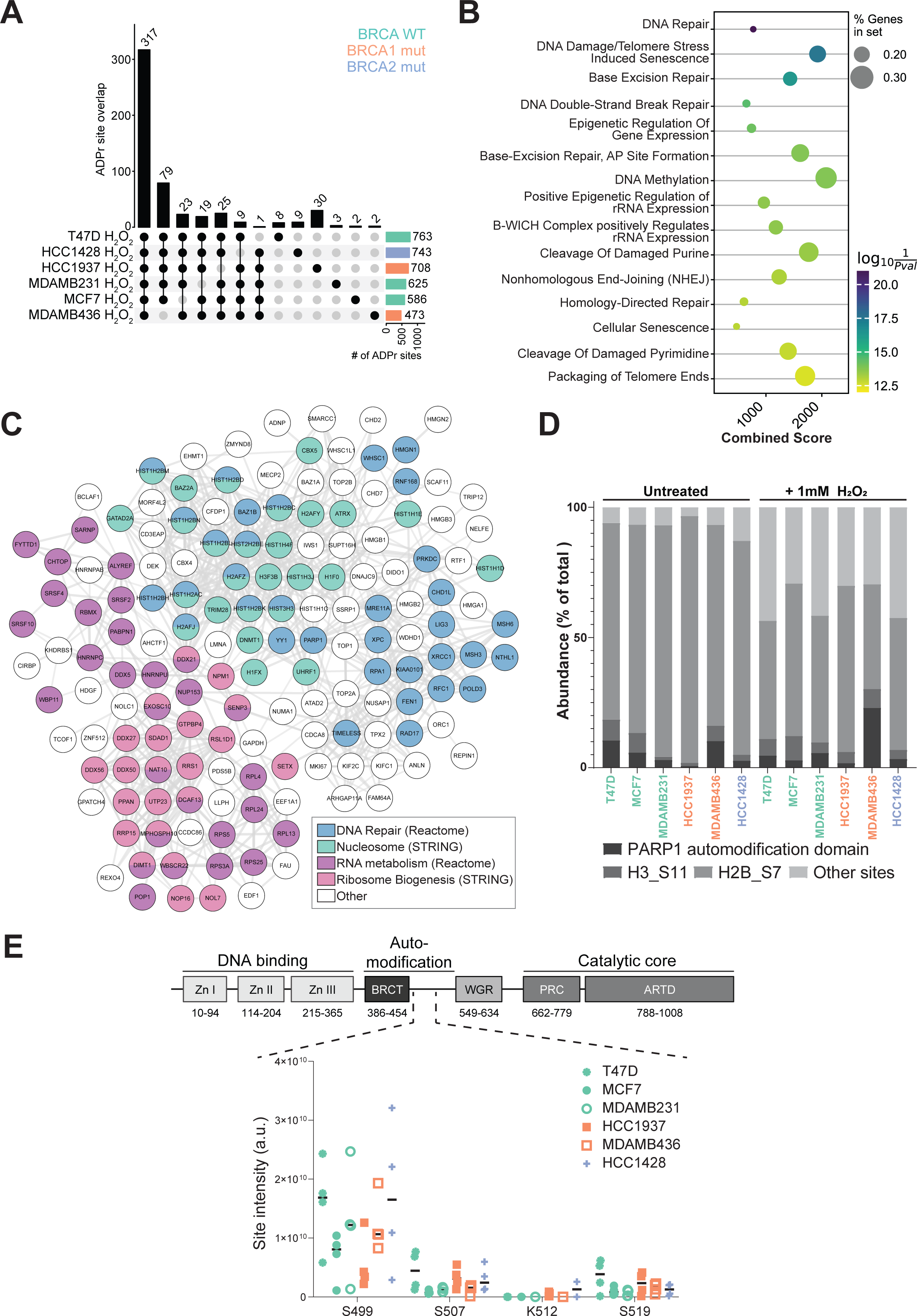
Serine ADPr of DNA damage repair proteins is induced in all cell lines in response to H_2_O_2_ treatment, with distinct differences in PARP inhibitor-resistant HCC1937 cells. **A.** UpSet plot showing the number of ADPr sites identified in all cell lines, 5 out of 6 cell lines, and exclusively in each cell line. **B.** GSEA was performed on the ADPr target proteins identified in all 6 cell lines, and the dot plot shows the top 15 enriched Reactome terms. The dot sizes represent the percentage of genes in the input set, and dots are colored according to log10 (1/P-value). The combined score is the log P-value calculated from a Fisher exact test multiplied by the z score of the deviation from the expected rank. GSEA were performed using Enrichr in GSEApy. **C.** The STRING network shows ADPr target proteins identified at the ADPr site level in all 6 cell lines. Proteins involved in DNA Repair, Nucleosome, RNA metabolism and Ribosome Biogenesis were highlighted in the indicated colors and the rest are gray. Proteins not connected to any others are excluded from the network. **D**. Intensity distribution of the three most abundant ADPr sites compared to all quantified sites in untreated and H_2_O_2_-treated cells as a percentage of total intensity. **E.** ADPr intensities for the PARP1 sites S499, S507, and S519 on the auto-modification domain after H_2_O_2_ treatment. PARP1 domain schematic was made with biorender.com.

Previous studies have highlighted histones as significant targets of ADPr^53-57^, with the PARP1 automodification domain also strongly targeted^58-60^. To explore whether this is similar in breast cancer cell lines, we compared the LFQ intensities of individual sites in each cell line to the total ADPr intensity in each cell line. We observed that in all cell lines, serine 6 (S6) on Histone H2B (hereafter referred to as H2BS7 for consistency with UniProt nomenclature) was the most abundantly ADP-ribosylated site both in untreated and H_2_O_2_-treated cells, followed by the PARP1 automodification domain (S499, S507, and S519), and S10 on Histone H3 (referred to here as H3S11) (**Fig. 4D**). Together, ADPr of these residues account for 85 – 90% and 55-75% of the ADPr intensity in untreated and H_2_O_2_-treated cells, respectively. In the PARP1 automodification domain, S499 was the most abundant ADPr site (**Fig. 4E**), with PARP1 automodification levels generally observed to be lowest in HCC1937 cells and highest in MDAMB436 cells, respectively. We also observed that histones H2B, H1, H3, PARP1, and HNRNPU were the top five ADP-ribosylated proteins in most of the cell lines by intensity (**Table 3**). Thus, despite variations in abundance across the breast cancer cell lines, histones and the PARP1 auto-modification domain are major ADPr targets, demonstrating a strong homogenous ADPr response to H_2_O_2_ triggered DNA damage in the cell lines, regardless of whether the cells have BRCA1 or BRCA2 mutations or are sensitive to PARP inhibitors.

### H_2_O_2_-stimulated ADP-ribosylation is conserved in mammalian and non-mammalian cells

Next, we compared the ADPr response after H_2_O_2_ treatment in the six breast cancer cell lines to other model systems and found that 1324 out of the 1632 sites identified across the breast cancer cells have previously been identified in HeLa cells after H_2_O_2_ treatment (**Fig. 5A**). On the protein level, 699 out of 777 proteins ADP-ribosylated in the breast cancer cell lines have been shown to also be ADP-ribosylated in HeLa cells upon H_2_O_2_ treatment^35^, additionally demonstrating the homogeneity of the ADPr response across different human cell lines. We also compared the ADPr sites identified in our dataset to those identified in *Drosophila* S2R+ cells^36^, because about 65% of human disease-causing genes have functional homologues in *Drosophila^61^.* We used flybase ID to find human orthologues for 219 for the proteins ADP-ribosylated upon H_2_O_2_ treatment in *Drosophila* proteins. Among those, 91 overlapped with ADPr target proteins identified in this study (**Fig. 5B**). We performed GSEA on the proteins found in both cell studies and observed that rRNA processing, RNA transcription, and translation pathways were enriched in both datasets (**Supp. Fig. 4A**), whereas the ADPr target proteins found exclusively in breast cancer cells are predominantly involved in DNA repair pathways (**Fig. 5C**) and RNA processing pathways are enriched among ADPr targets in *Drosophila* (**Supp. Fig. 4B**).

**Figure 5.**
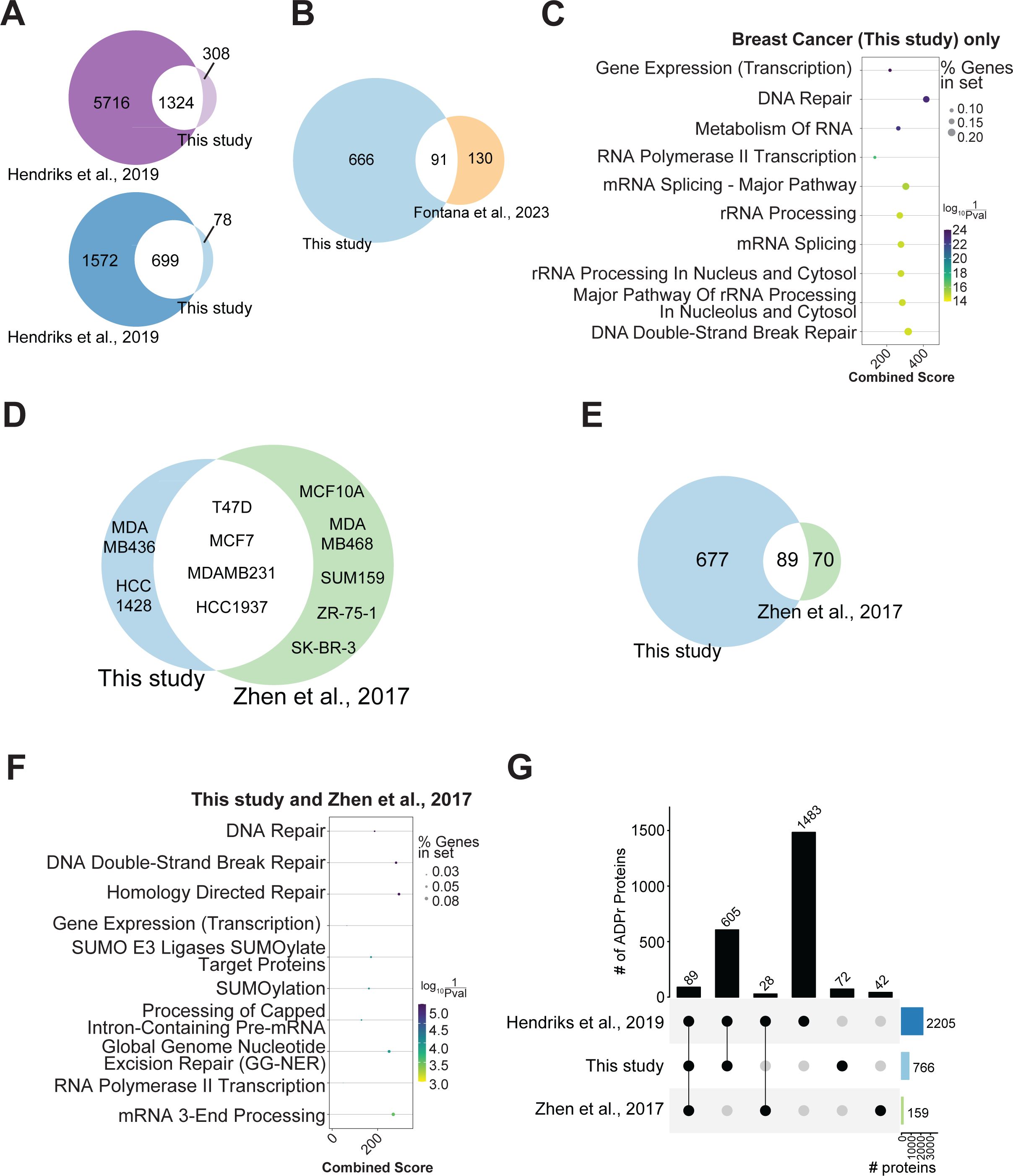
H_2_O_2_-stimulated DNA damage ADPr is conserved in mammalian and non-mammalian cells. **A.** Venn diagrams showing the site-level (top) and protein-level (bottom) overlap in ADPr targets identified in breast cancer cells (this study) and HeLa cells (Hendriks et al., 2019). **B.** Venn diagram showing the overlap between the ADPr target proteins identified in this study and matching human orthologues of ADPr target proteins identified in *Drosophila* S2R+ cells. **C.** GSEA was performed on the 666 proteins from (B) identified exclusively in breast cancer cells, and the top 10 enriched Reactome pathways were plotted on a dot plot. **D.** Venn diagram showing the breast cancer cell lines used in this study and in the study by Zhen et al., profiling D/E-ADPr in H_2_O_2_-treated breast cancer cells. **E.** Venn diagram showing the protein-level overlap in the four breast cancer cell lines common to this study and the D/E-ADPr profiling study. **F.** GSEA performed on the proteins from (**E**) for the 89 ADPr target proteins identified in this study and the Zhen et al. study, with the top 10 enriched Reactome pathways plotted on a dot plot. In C and G, the dot sizes represent the percentage of genes in the input set, and dots are colored according to log10 (1/P-value). The combined score is the log P-value calculated from a Fisher exact test multiplied by the z-score of the deviation from the expected rank. GSEA were performed using Enrichr in GSEApy. **G.** UpSet plot showing the protein-level overlap between the predominantly serine-ADPr target proteins (this study), D/E-ADPr target proteins in the four common cell lines (Zhen et al., 2017), and the predominantly serine-ADPr target proteins in HeLa cells (Hendriks et al., 2019).

We further wanted to compare our breast cancer H_2_O_2_-stimulated ADPr sites to another study looking at ADPr in breast cancer cell lines^31^. This study, where a method to profile ADPr on D and E residues was employed^62^, reported that DNA damage-induced D/E-ADPr was very heterogeneous and highly cell line-specific in breast cancer cells^31^. Given the predominance of serine ADPr sites in our study, a direct site-specific comparison to D/E-ADPr was not feasible. Instead, we compared the proteins targeted by ADPr across the four cell lines used in both studies (T47D, MCF7, MDAMB231, and HCC1937) (**Fig. 5D**). Collectively, we found that 89 of the 159 D/E-ADPr proteins were also identified in our predominantly serine-ADP-ribosylome (**Fig. 5E**). GSEA revealed that the common target proteins were involved in various DNA repair pathways, including double-strand break repair, homology-directed repair, and base excision repair, underscoring the importance of ADPr in these processes (**Fig. 5F, Supp. Fig. 4C&4D**). We furthermore observed that the 89 ADPr proteins common to this study and in the D/E-ADP-ribosylome were also identified in HeLa cells, with 28 additional proteins in the D/E-ADP-ribosylome also identified as serine ADPr targets in HeLa cells (**Fig. 5G**). Together, these results demonstrate a considerable overlap of protein involved in DNA repair and RNA processing targeted by ADPr after H_2_O_2_ treatment, both in mammalian and non-mammalian cells.

### ADP-ribosylation of the tumor suppressor USF1 is significantly up-regulated in PARPi sensitive cell lines

Considering the remarkable overlap between sites targeted for serine ADPr after H_2_O_2_-induced DNA damage across the six breast cancer cell lines and other cell lines, we wondered if there would be differences in the abundances of these sites between PARPi sensitive and resistant cell lines. To investigate this, we first compared the LFQ intensities of the 899 sites quantified among the six breast cancer cell lines between PARPi sensitive cells (MDAMB436 and HCC1428) versus BRCA wild type cells (T47D, MCF7, and MDAMB231), as well as PARPi sensitive cell lines versus the PARPi resistant cell lines (T47D, MCF7, MDAMB231, and HCC1937). We decided to use this very stringent approach, since this would allow us to look beyond differences arising from the molecular subtypes of the different cell lines and highlight ADPr regulation specific to PARPi sensitive and resistant cells. As expected, only six different ADPr sites were significantly different in abundance between the groups in both comparisons (**Fig. 6A&6C**). Furthermore, most of these regulated ADPr sites resided on proteins, which were also expressed differently between the two groups (**Fig. 6B&6D**). Interestingly, there was one ADPr event, which was consistently significantly down-regulated in the BRCA WT and PARPi resistant cell lines compared to the PARPi sensitive cell lines – S189 on Upstream stimulatory factor 1 (USF1) (**Fig. 6A&6C**), whereas its protein expression was unchanged (**Fig. 6B&6D**). USF1 is a transcription factor and a stress responsive tumor suppressor, and its activity has been shown to be regulated by PTMs **(Fig. 6E**)^63-65^.

**Figure 6.**
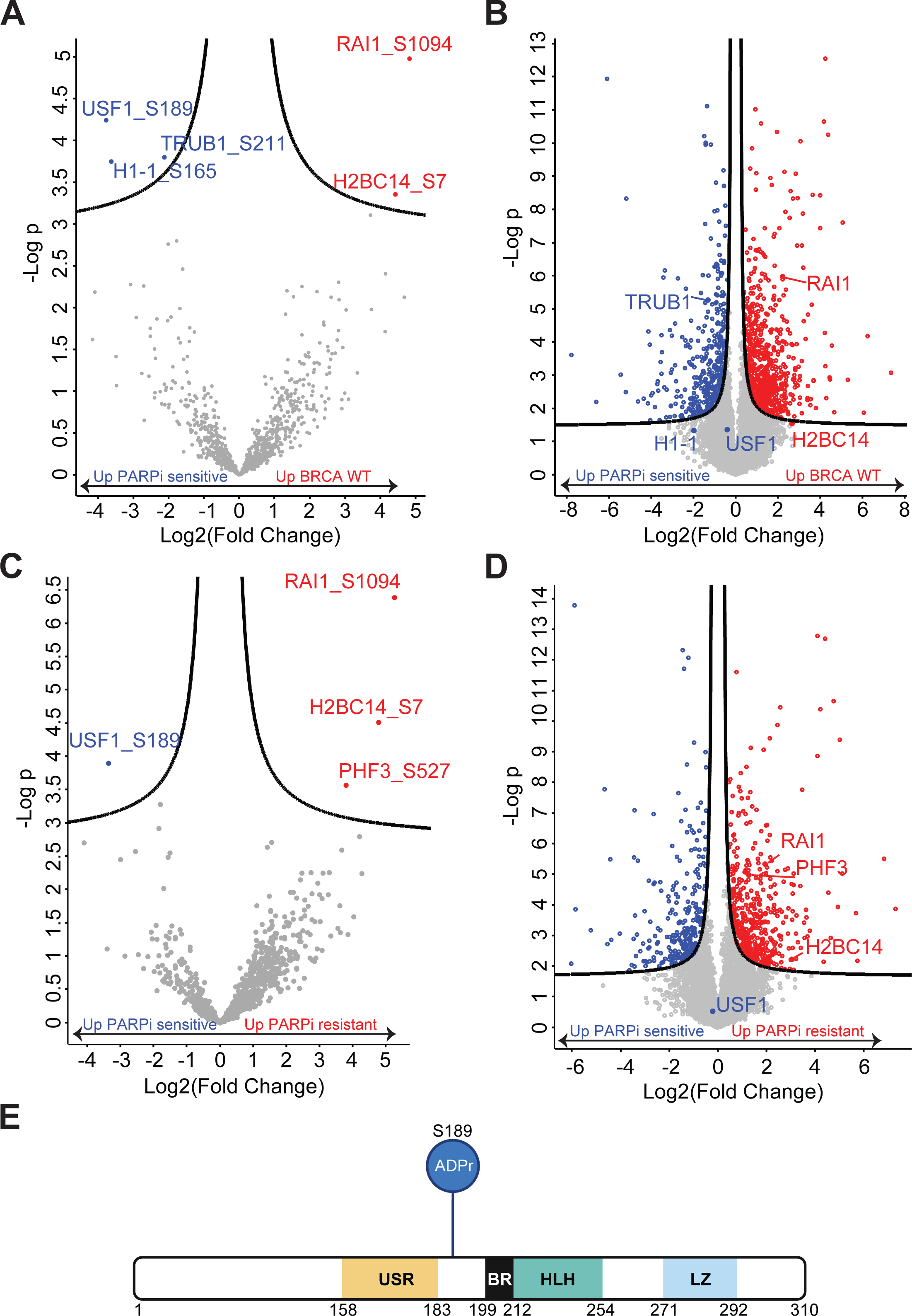
USF1 is significantly upregulated at the ADPr site level in PARPi resistant breast cancer cells. **A.** The volcano plot shows ADPr sites significantly up (red) or down (blue) in PARPi-sensitive cells (MDAMB436 and HCC1428) compared to BRCA WT cells (T47D, MCF7 and MDAMB231) at a False Discovery Rate (FDR) of 0.1. **B.** The volcano plot shows proteins significantly upregulated (blue) or downregulated (red) in PARPi-sensitive cells compared to BRCA WT cells (FDR = 0.01), detected by MS proteome profiling described in Figure 2. **C.** The volcano plot shows ADPr sites significantly up (red) or down (blue) in all PARPi-resistant (T47D, MCF7, MDAMB231, HCC1937) compared to PARPi-sensitive (MDAMB436 and HCC1428) cells at 0.1 FDR. **D**. The volcano plot shows proteins significantly upregulated (blue) or downregulated (red) in PARPi-sensitive cells compared to PARPi-resistant cells (FDR = 0.01), detected by MS proteome profiling described in Figure 2. Proteins with significantly upregulated or downregulated ADPr sites are labeled. Volcano plot p-values are based on two-sided T-tests with multiple testing correction. The list of significantly regulated ADPr sites and statistics are in **Supplemental Table 3**. **E.** Schematic of USF1 domains with ADPr on S189 highlighted. Domain organization is given according to Sirito et al^88^. USR – USF-specific region; BR – basic region; HLH – helix-loop-helix; LZ – leucine zipper. Schematic was made with biorender.com.

### The PARPi-resistant HCC1937 cell line has several unique ADPr target proteins and longer PAR chains

Overall, there were very few quantified sites specific to one cell line or specific to PARPi sensitive or resistant cell lines (**Supp. Fig. 3B & 3C and Table 4**). Notably, despite the homogeneity of serine ADPr across the cells, the PARPi resistant BRCA1 mutant cell line HCC1937 had the highest number (30) of quantified ADPr sites not found in any other cell lines (**Fig. 4A**). These 30 sites were found on 23 proteins (**Fig. 7A**), 11 of which were ADP-ribosylated on other acceptor sites in one of more of the other cell lines. The remaining 12 proteins exclusively ADP-ribosylated in HCC1937 cells include PARP2, which is known to exert functional redundancy with PARP1 in DNA damage repair^66^. Despite their unique ADPr status, most of the 23 proteins were expressed in the other cell lines, suggesting that the specific lack of ADPr in these cell lines is not due to lack of protein expression (**Fig. 7B**). Of note, neither PARP2 nor GOLGA6A were detected in any cell lines in our proteome dataset, indicating an overall low expression of these proteins (**Supp. Table 1**).

**Figure 7.**
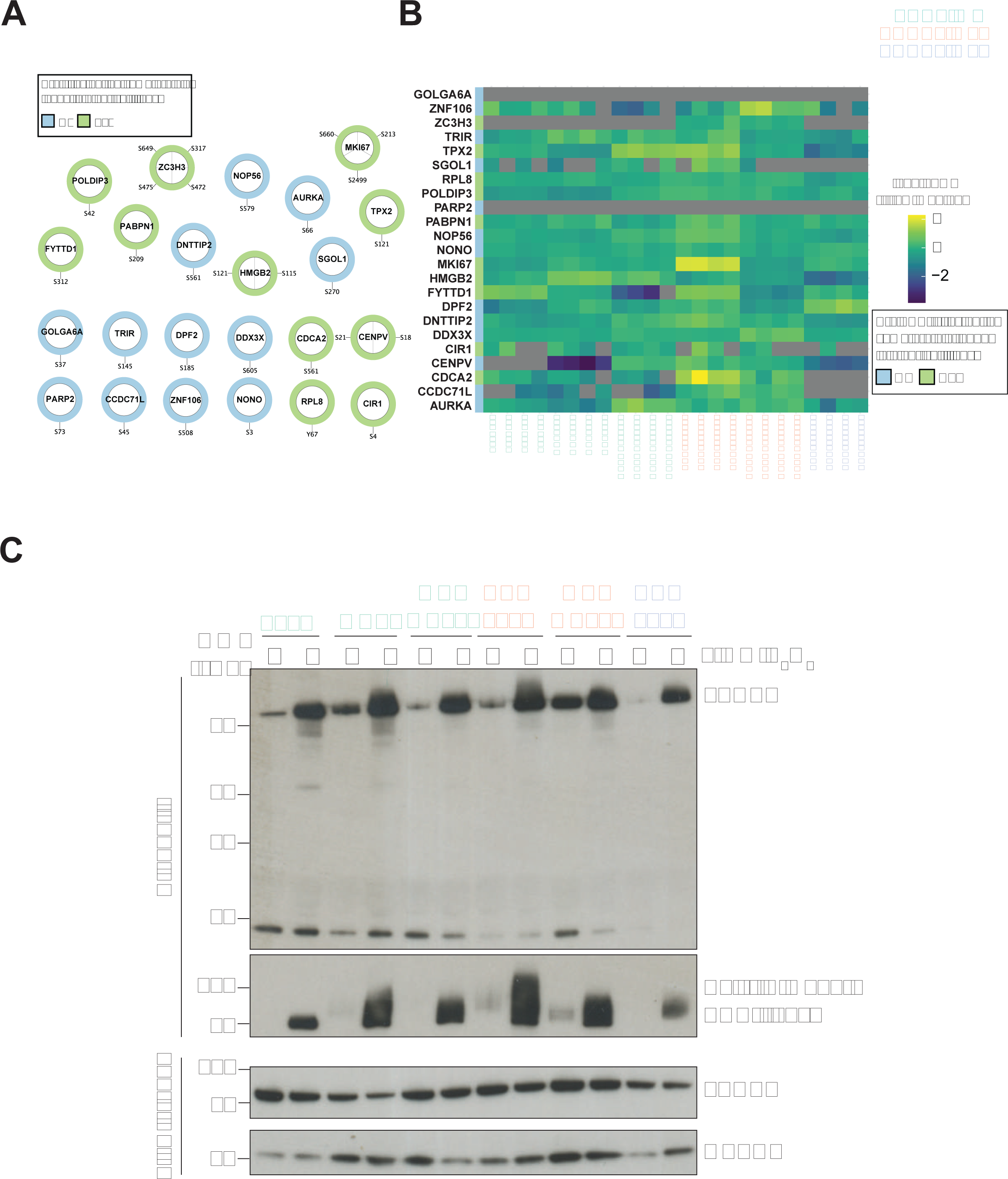
PARPi resistant BRCA1 mutant HCC1937 cells has several distinct H_2_O_2_-stimulated ADPr targets and longer PAR chains. **A.** STRING network showing ADPr sites identified exclusively in HCC1937 cells. **B.** Heatmap showing the protein levels of HCC1937-exclusive ADPr targets in all cell lines, detected by mass spectrometry-based proteome profiling described in Figure 2. **C.** Western blot showing Poly-ADPr chains on PARP1 and PARylated PARP1 levels in Af1521 affinity-purified proteins from untreated or H_2_O_2_-treated cell lines.

Considering the differences we observe in both the PARPi sensitivity of and ADPr sites identified in HCC1937 cells compared to the other BRCA mutants, we wanted to further explore the ADPr response in HCC1937 cell lines compared to the other cell lines. As our proteome data shows a much lower expression levels of the PARG glycohydrolase, which digests PAR chains into MAR (**Fig. 1B & Fig. 2E**), we assessed the length of ADPr chains in this cell line. To this end, we treated cells with 1 mM H_2_O_2_ for ten minutes, affinity purified ADP-ribosylated proteins using the Af1521 domain^67^, and subsequently stained for PARP1 along with PAR and MAR by WB. Similar amounts of PARP1 were pulled down in HCC1937 cells compared to other cell lines, indicating that PARP1 is ADP-ribosylated to a similar degree in these cell lines (**Fig. 7C**). However, PAR chains on PARP1 were longest in the ‘PARG-low’ HCC1937 cells. In general, we observe that lower levels of PARG (**Fig. 1B**) are associated with increased PAR chain length in the cell lines. This provides further evidence that PARP1 expression and activation is homogeneous across the breast cancer cells and suggests that HCC1937 cells have longer ADPr chains because PARG expression is decreased. Overall, these results show that although H_2_O_2_ treatment induces a strong and homogeneous DNA damage response in all the breast cancer cells investigated in this study, there are several unique ADPr protein targets in PARPi resistant BRCA mutant HCC1937 cells. Moreover, we find that the observed low level of PARG expression in this cell line leads to longer PAR chains, which might have implications for the DNA damage response and PARPi sensitivity in this cell line.

## Discussion

In this study, we profiled DNA damage ADPr in a panel of six breast cancer cell lines with differences in BRCA mutation status and PARPi sensitivity using MS-based proteomics. We identified 1632 and quantified 899 ADPr sites in total and found that 92.8% of those target serine residues. We found that a large part of the ADP-ribosylation sites in PARPi sensitive and PARPi resistant breast cancer cells overlap. Furthermore, comparison with previously published reports identifying proteome-wide ADPr target sites after H_2_O_2_ treatment^35,36^ demonstrates high site- and protein-level overlap with other mammalian and non-mammalian cell types. This supports to the idea of a highly conserved and homogeneous core ADPr signaling network in response to H_2_O_2_ induced DNA damage.

The homogeneity of the predominantly serine-ADPr response to H_2_O_2_-induced DNA damage in our study is striking in contrast to the heterogeneity in glutamic acid- or aspartic acid-linked ADP-ribosylation in the same breast cancer cell lines from an earlier publication^31^. This previous study used chemical modification of D/E-ADPr followed by enrichment for peptides carrying these chemically modified sites^62^. The specific enrichment technique, while limiting in other aspects, could allow for identification of low abundant D/E-ADPr sites. In our study, however, ADP-ribosylated peptides were directly enriched using an Af1521 macrodomain, which specifically binds ADPr moieties^68^. MS-analysis in our study was carried out with EThcD fragmentation, a technique gentle enough to keep the labile ADPr group intact while fragmenting the peptide backbone^69^, enabling accurate site-level identification of any ADP-ribosylated amino acid without the need for derivation^33,70^ and allowing us to profile ADPr sites targeting a wider range of amino acids. Another major difference between the two studies is that in the D/E-ADPr characterization study, stable PARG depletion was performed using shRNAs prior to H_2_O_2_ treatment and D/E-ADPr profiling^62^, which could introduce differences in the accumulation of ADPr sites in the cells at the time when samples are harvested. In contrast, our method only inhibits PARG activity during cell lysis with denaturing buffers immediately after H_2_O_2_ treatment. However, we do not exclude that sample preparation conditions, low abundance of the sample measured in the MS, and the stochastic nature of DDA MS could preclude the identification of other amino acids to similar depth as serine. Despite these differences, we still see a large overlap between the proteins identified as ADPr targets in both studies, as well as with ADPr sites identified in HeLa cells, emphasizing the importance of ADP-ribosylation of these proteins in response to DNA damage.

In this as well as in the other studies discussed in this paper, histones are heavily targeted by ADPr, and comprise a large percentage of the total ADPr signal. Because ADPr is a bulky molecule with two negative charges at physiological pH, ADPr of a protein can alter its properties and functions^71^. Here, we found that histones H2BS7 and H3S11 are major ADPr targets in all the breast cancer cell lines, and ADPr of H2BS7 and H3S11 has been shown to convert nucleosomes into robust substrates for the chromatin remodeler ALC1^72^. Moreover, H3S11 is phosphorylated by Aurora B kinase during mitosis^73^, and this is counteracted at UV-damaged chromatin by ADPr^13,74^. ADPr of histone H3S11 is also known to be mutually exclusive with H3K10 (Uniprot numbering) acetylation^33,75,76^, raising the possibility for PTM crosstalk in breast cancer cells. Although we demonstrate that DNA damage induced ADPr is homogenous at the site level and conserved between various cell lines, we did find notable differences in the abundances of individual modification sites between PARPi resistant and sensitive cell lines. Strikingly, when comparing H_2_O_2_-induced ADPr in all PARPi sensitive cell lines to the PARPi resistant breast cancer cell lines, we found that the ADPr on S189 in the protein USF1 was significantly down-regulated in all the PARPi resistant cell lines – including the PARPi resistant BRCA mutant HCC1937. USF1 is a stress responsive transcription factor and tumor suppressor gene. It can inhibit cell growth, proliferation, cell cycle progression, and prevent transformation of primary cells by c-Myc, and loss of function of USF1 has been shown to favor proliferation in transformed breast cancer cell lines^63,65^. Furthermore, previous studies have shown that the activity of USF1 can be regulated by PTMs^64^, whereas here we find down-regulation of the addition of the bulky ADPr modification to this protein specifically in PARPi resistant breast cancer cell lines, raising questions about the implications of this modification on the activity of USF1. It is tempting to speculate that the upregulation of ADPr on S189 of USF1 could change the activity of this tumor suppressor and have implications for the cell proliferation and survival of the PARPi sensitive cell lines studied here.

Finally, we found very different levels of PARPi sensitivity in the three tested BRCA mutant cell lines, with BRCA1 mutant MDAMB436 being most PARPi sensitive whereas BRCA1 mutant HCC1937 cells were not sensitive to any of the three PARPi tested. In MDAMB436 cells, we observed that ADPr PARP1 is highest in both untreated and H_2_O_2_ treatment conditions compared to the other cells. On the protein level, ARH3 expression is also highest in these cells, suggesting that MDAMB436 cells may be more reliant on PARP1-mediated ADPr signaling for DNA repair through alternative pathways. This increased PARP activity, however, presents a therapeutic vulnerability, as MDAMB436 cells are most sensitive to PARPi. Interestingly, in the PARPi resistant HCC1937 cell line, we observed that PARG levels were decreased compared to the other cell lines. We also found that as a result of this, HCC1937 cells had longer PAR chains compared to the other five breast cancer cell lines and the most unique ADPr sites in H_2_O_2_-treated conditions. Together, these observations suggest that by downregulating PARG expression, DNA damage-induced PARP1 activation persists for longer. The cells may therefore be able to tolerate higher PARPi doses as we observed, even when they lack functional BRCA1. In fact, downregulation of PARG was shown to restore PAR formation and partially rescue PARP1 signaling, counteracting the synthetic lethality of PARP inhibitors in BRCA2-mutated mouse tumors^77^.

In summary, by profiling the ADP-ribosylomes of six breast cancer cells and comparing them to other datasets, we have demonstrated that the ADPr response to DNA damage is much more homogenous and robust across various cell systems compared to what has been thought before. Despite this, we have identified unique differences between PARPi sensitive and resistant cell lines, which could inform future studies aiming to better understand the development of PARPi resistant in breast cancer.

## Materials and methods

### Cell lines

T47D, MCF7, MDAMB231, HCC1937, MDAMB436, and HCC1428 cells, originally from the American Type Culture Collection, were kindly provided by Jesper Olsen’s lab (NNF-CPR, University of Copenhagen). Cells were propagated in Dulbecco’s modified Eagle’s medium (Invitrogen) supplemented with 10% fetal bovine serum and penicillin/streptomycin (100 U/mL, Gibco) (cell culture medium) and incubated at 37°C with 5% CO_2_. Cells were routinely tested for mycoplasma contamination. When not in culture, cells were stored in fetal bovine serum with 5% DMSO in a liquid nitrogen freezer.

### HRD score and HRDetect score predictions

#### Genomic scar scores

Collectively, the sum of three individual scores, consisting of the HRD-LOH, HRD-LST and HRD-TAI is termed the HRD score^37^. The HRD-LOH score quantifies the number of instances where LOH (loss of heterozygosity) regions surpass 15 Mb in length without encompassing the entirety of the chromosome^78^. The Large Scale Transitions (HRD-LST) score is defined by the count of chromosomal breaks between proximate regions spanning at least 10 Mb, while the intervening space does not exceed 3 Mb^79^. The number of Telomeric Allelic Imbalances (HRD-TAI) measures the instances where allelic imbalances (AIs)—defined as a disproportionate representation of parental allele sequences—extend to a chromosome’s telomeric region^80^. The HRD score was calculated using the scarHRD R package^81^, using whole exome sequences of breast cancer cell lines downloaded from the Cancer Cell Line Encyclopedia (CCLE). A cut-off value of >= 42 was used following the recommendations by How et al^82^.

#### HRDetect score

The HRDetect score for the breast cancer cell line samples were calculated using the weights of a previously published model, which was trained on a dataset of 560 synthetic whole exomes from the TCGA (The Cancer Genome Atlas) breast cancer cohort^83^. The same cut-off value (>=0.7) was used as defined with the original HRDetect score.

#### Western blot

Cell pellets were lysed in a denaturing SDS lysis buffer (50 mM Tris pH 8.5, 150 mM NaCl and 2% SDS) and homogenized by incubating at 90°C with shaking at 1400 rpm for 1 hour on a benchtop heating block. Protein concentrations were quantified by BCA (Thermo Fisher) according to the manufacturer’s protocol. NuPAGE LDS sample buffer (Invitrogen) was added to cell lysates to a final concentration of 1X, and lysates were boiled at 95°C for 5 minutes. To blot for BRCA2, lysates were separated by sodium dodecyl sulfate/polyacrylamide (SDS/PAGE) gel electrophoresis on 1 mm NuPAGE 3 – 8% gradient Tris-Acetate gels (Thermo Fisher) using Tris-Acetate SDS running buffer and transferred to nitrocellulose membranes (Bio-Rad) with NuPAGE transfer buffer supplemented with 10% methanol using a Trans-Blot Turbo Transfer system (Bio-Rad). For all other antibodies, lysates were separated by SDS-PAGE gel electrophoresis on 1 mM NuPAGE 4 – 12% bis-tris gels (Thermo Fisher) using MOPS running buffer and transferred to nitrocellulose membranes using Tris-Glycine-methanol transfer buffer (30.3 g Tris Base, 144g Glycine, and 20% Methanol in 1000 mL). Protein loading amounts were assessed with Ponceau staining, after which the membranes were washed for 10 minutes in PBS supplemented with 0.1% Tween-20 (PBS-T). Membranes were blocked for 1 hour using 5% BSA or 5% milk in PBS-T depending on the primary antibody used. Membranes were then incubated with primary antibodies overnight at 4°C and washed three times with PBS-T. Membranes were then incubated with HRP-conjugated secondary goat antibodies to mouse or rabbit diluted 1:7500 in PBST with 5% milk (Rockland) for 1 hour at room temperature, washed three times with PBST, and proteins were detected using the Novex ECL Chemiluminescent Substrate Reagent Kit (Invitrogen). The following antibodies were used in this study: PARP1/2 Rabbit pAB sc-7150 (Santa Cruz Biotech) 1:200, PARG (D4E6X) Rabbit mAb #66564 (Cell Signaling), HPF1 Rabbit pAb HPA043467 (Atlas Antibodies), ARH3/ADPRHL2 Rabbit pAb HPA027104 (Atlas Antibodies), BRCA1 Rb mAb sc-6954 (Santa Cruz Biotech), BRCA2 Rb mAb #10741 (Cell Signaling), MSH2 (D24B5) XP Rb mAb #2017 (Cell Signaling), MSH6 (D60G2) XP Rb #5424 (Cell Signaling), GAPDH Rabbit pAb ab9485 (Abcam) 1:2000, gamma-Tubulin Ms mAb T5326 (Sigma Aldrich) 1:2500. PARP1, PARG, HPF1, ARH3, BRCA1, and gamma-Tubulin antibodies were diluted in PBST with 5% BSA, and BRCA2, Poly/Mono-ADP Ribose, GAPDH, MSH2, and MSH6 antibodies were diluted in PBST with 5% milk. Antibodies were used at a dilution of 1:1000 unless otherwise indicated.

#### Clonogenic survival assays

Olaparib (AZD2281 Ku-0059436), Rucaparib phosphate (AG-0144498 PF-01367338), and Talazoparib (BMN673) (all from Selleck Chemicals) were dissolved to 10 mM stocks in DMSO. Cells were seeded sparsely and evenly at 1000/well (T47D, MCF7, HCC1937, MDAMB436, and HCC1428) or 400/well in 6-well plates and incubated overnight. The next day cells were treated with 0.00093, 0.0056, 0.033, 0.2, 1, 5, or 10 µM Olaparib, 0.0067, 0.033, 0.167, 1, 2, 5, or 10 µM Rucaparib, or 0.000043, 0.00026, 0.0015, 0.0093, 0.056, 0.33, or 2 µM Talazoparib diluted in cell culture medium with a final DMSO concentration of 0.1%. Cell culture medium with 0.1% DMSO was added to control wells, and each treatment was done in triplicate. Cells were continuously exposed to drug for 9 – 18 days, and cell culture medium containing the drugs was refreshed every 3 – 4 days. After incubation, cells were fixed and stained with 0.5% crystal violet staining solution with 20% Methanol. Colony survival was calculated as a percentage relative to DMSO-treated conditions.

#### Sample preparation for proteome profiling

For proteome mass spectrometry experiments, breast cancer cells were grown to 80% confluency, washed twice with cold PBS, harvested on ice with a cell scraper, and pelleted by centrifuging at 400 x *g* for 3 minutes and aspirating off PBS. Cells were lysed with 2mL chaotropic denaturing GndHCl lysis buffer (6M guanidine-HCl (G3272; Sigma Aldrich), 50 mM Tris pH 8.5). 5 mM tris(2-carboxyethyl)phosphine (TCEP; C4706, Sigma Aldrich) and 10 mM chloroacetamide (CAA; 22790, Sigma Aldrich) were added to lysates for reduction of disulfide bonds and methionine alkylation, respectively. To shear DNA, lysates were sonicated on ice for 45 seconds at 60% amplitude. After 1 hour incubation at room temperature, proteins were digested into peptides by incubating with Lys-C endopeptidase (1:200 w/w; Wako Chemicals) for 3 hours, followed by overnight incubation with modified sequencing-grade Trypsin (1:200 w/w; Sigma Aldrich) after diluting to 1.5M GndHCl with 25 mM Tris pH 8.5. Trifluoroacetic acid (TFA) was added to a final concentration of 0.5% v/v to inactivate proteases, and samples were centrifuged at high speed to remove precipitates. Peptides were purified using reverse-phase Sep-Pak C18 cartridges (WAT051910; Waters) according to manufacturer’s instructions, eluted in 30% Acetonitrile (ACN) in 0.1% TFA, and vacuum-dried in in a SpeedVac (Eppendorf) at 60 °C. Purified peptides were reconstituted in 50 mM Ammonium bicarbonate (ABC) and separated on an Acuity UPLC Peptide CSH C18 1.7 µm reversed-phase column (186006935; Waters) under basic conditions, with fractions collected in 46 timed intervals and concatenated into 12 fractions (Batth and Olsen, 2016). Formic acid was added to peptides to a final concentration of 0.5% v/v and peptides were vacuum-dried at 60 °C. Peptides were reconstituted in 0.1% Formic acid for mass spectrometry analysis.

#### Cell culture, lysis, and enrichment of ADP-ribosylated peptides

For ADPr mass spectrometry experiments, breast cancer cells grown to 80% confluency were treated with 1 mM H_2_O_2_ in PBS for 10 minutes at 37 °C and washed once with PBS at 4 °C. Untreated cells were washed twice at 4 °C with cold PBS. Cells were collected by gentle scraping and pelleted by centrifugation at 400 x *g* for 3 minutes at 4 °C. ADP-ribosylated peptides were enriched as previously described^35,49,50^. Briefly, cell pellets were lysed by adding 10 pellet volumes of guanidinium-HCl lysis buffer, alternating vigorous shaking and vortexing, and finally snap freezing in liquid nitrogen. Lysates were brought to a thaw at room temperature, treated with 5 mM TCEP (C4706, Sigma Aldrich) and 5 mM CAA (22790, Sigma Aldrich) for alkylation and reduction, and sonicated on ice for 45 seconds at 60% amplitude. Proteins were digested by incubating with Lys-C endopeptidase (1:200 w/w; Wako Chemicals) for 3 hours, followed by overnight incubation with modified sequencing-grade Trypsin (1:200 w/w; Sigma Aldrich) after diluting to 1.5M guanidine-HCl with 50 mM ABC. TFA was added to a final concentration of 0.5% v/v to inactivate proteases and samples were centrifuged at high speed to remove precipitates. Peptides were purified using reverse-phase Sep-Pak C18 cartridges (WAT051910; Waters) according to manufacturer’s instructions, eluted in 30% ACN in 0.1% TFA, frozen at least overnight at -80 °C, and lyophilized for 96 hours. Peptides were reconstituted in AP buffer (50 mM Tris-HCl (pH 8.0), 50 mM NaCl, 1 mM MgCl2, and 250 µM DTT) and 3 mg of each sample was aliquoted. Recombinant hPARG (1:10,000 w/w, a kind gift from Prof. Michael Höttiger) was added to 3 mg of each sample and incubated overnight at room temperature with gentle shaking to reduce ADP-ribose polymers, and precipitates were removed by centrifuging at 4 °C for 30 min at 4250 × *g*. GST-tagged Af1521 macrodomain beads were produced in-house using BL21(DE3) bacteria and coupled to glutathione Sepharose 4B beads (Sigma-Aldrich), essentially as previously described. Peptides were incubated with GST-tagged Af1521 macrodomain beads (100 µL dry beads per 10 mg sample) and incubated in a head-over-tail mixer at 4 °C for 3 hours. Beads were washed twice with ice-cold AP buffer, twice with ice-cold PBS with 250 µM DTT, and twice with ice-cold water. ADP-ribosylated peptides were eluted with 0.15% TFA and centrifuged through 0.45 µM spin filters, followed by centrifugation through 100 kDa cut-off spin filters (Vivacon). Peptides were purified and fractionated on stage tips at high pH into 2 fractions. After loading onto stage tips for high pH fractionations, the flowthrough was collected and subjected to low pH stage tip purification as previously described.

#### Mass spectrometric analysis

Proteome mass spectrometry experiments were performed on an Orbitrap Exploris 480 mass spectrometer (MS) using high-energy collisional dissociation (HCD) fragmentation. ADPr mass spectrometry experiments were performed on an Orbitrap Fusion Lumos™ Tribrid™ MS (Thermo) using electron-transfer/higher-energy collision dissociation (EThcD) fragmentation. Samples were analyzed on 15 – 20-cm long analytical columns with an internal diameter of 75 µm, packed in-house using ReproSil-Pur 120 C18-AQ 1.9 µm beads (Dr. Maisch) connected to a nanoscale EASY-nLC 1200 liquid chromatograph (Thermo). The analytical column was heated to 40 °C using a column oven and peptides were eluted from the column with a gradient of buffer A (0.1% formic acid) and buffer B (80% ACN in 0.1% formic acid). For proteomes, the primary gradient ranged from 5% to 35% buffer B over the course of 50 minutes, followed by an increase to 90% buffer B over 4 minutes, constant 90% buffer B for 2 minutes, decrease to 5% buffer B over 2 minutes, and 5% buffer B for 2 minutes. For high pH fractions of ADPr samples, the primary gradient ranged from 3% to 38% buffer B over the course of 38 minutes, followed by an increase to 90% buffer B over 2 minutes, constant 90% buffer B for 4 minutes, decrease to 5% buffer B over 3 minutes, and 5% buffer B for 3 minutes. For low pH fractions of ADPr samples, the primary gradient ranged from 5% to 15% buffer B over the course of 22 minutes, followed by an increase to 30% buffer B over 6 minutes, 90% buffer B over 3 minutes, constant 90% buffer B for 3 minutes, decrease to 5% buffer B over 3 minutes, and 5% buffer B for 3 minutes. Electrospray ionization (ESI) was achieved using a NanoSpray Flex NG ion source (Thermo). Spray voltage was set to 2 kV, capillary temperature to 275 °C, and RF level to 40%. Full scans were performed at a resolution of 120,000, with a scan range of 300–1750 m/z, and maximum injection time set to auto. The normalized AGC target was “200” for proteome and “150” for ADPr samples. For proteome samples, precursor isolation was performed at a width of 1.3 m/z, a normalized AGC target of “200”, and precursor fragmentation using HCD at 25% normalized collision energy (NCE). Top 18 precursors with charge state 2–6 were isolated for MS/MS analysis, and a dynamic exclusion of 60 s was used. MS/MS spectra were measured in the Orbitrap, with maximum precursor injection time set to auto and scan resolution of 15,000. For ADPr samples, precursor isolation was performed at a width of 1.3 m/z, normalized AGC target of “400”, and fragmentation using EThcD with an NCE of 20. Top 3 precursors with charge state 3–5 were isolated for MS/MS analysis, and a dynamic exclusion of 60 s was used. MS/MS spectra were measured in the Orbitrap, with a maximum precursor injection time of 1000 ms, and a scan resolution of 60,000.

#### Analysis of proteome data

The raw MS data was analyzed using MaxQuant software version 2.0.3.1 with default settings unless indicated. A human fasta file downloaded on 02.03.2022 from Uniprot.org was used to generate a theoretical spectral library. Label free quantification via Fast LFQ was enabled and normalization type was set to classic. Match between runs was enabled with a match time window of 0.7 minutes and alignment time window of 20 minutes (default parameters). From the list of proteins in the proteinGroups.txt file, the data was further filtered to remove potential contaminants, reverse hits, proteins only identified by site, and proteins quantified with LFQ intensity in fewer than three replicates of at least one cell line.

#### Analysis and filtering of ADP-ribosylome data

The raw MS data was analyzed using MaxQuant software version 2.1.3.0 with default settings unless indicated. A human fasta file downloaded on 11.03.2021 from Uniprot.org was used to generate a theoretical spectral library. The maximum missed cleavages was set to 6. Methionine oxidation, N-terminal acetylation, cysteine carbamidomethylation, and ADPr on cysteine, aspartic acid, glutamic acid, histidine, lysine, arginine, serine, threonine, and tyrosine residues were set as variable modifications, and the maximum number of modifications per peptide was set to 5. Label free quantification by fast LFQ was enabled and normalization type set to none. Match between runs was enabled with a match time window of 0.7 minutes and alignment time window of 20 minutes. Beyond the automatic filtering and FDR of 0.01 set by MaxQuant, the data was further manually filtered with the statistical software Perseus^84^ and Microsoft Excel to ensure proper identification and localization of ADPr sites as follows: potential contaminants, reverse hits, proteins only identified by site were removed, peptide-spectrum matches (PSMs) with more than one ADPr modification were excluded, and only ADPr site assignments with localization probability above 0.9 were included for site identification and above 0.75 for quantification. Because default MaxQuant intensity assignments to modification sites also include non-localized or poorly localized evidences, intensity values were manually mapped from the evidence.txt to the sites table based on localized PSMs only.

#### Comparisons to Zhen et al., 2017 characterization of D/E-ADP-ribosylation in breast cancer cells

Zhen and colleagues used the discontinued IPI accession numbers to identify their proteins, while we used Uniprot IDs (references). They also used gene names, but because some of the gene names were different (THOC/ALYREF, for example), we converted their gene names to Uniprot IDs and then compared them to our Uniprot IDs. For this reason, we did not consider hyphenated Uniprot IDs corresponding to isoforms of the same protein, and the number of proteins in our study is 505 instead of 511. The overlaps between ADPr proteins in this study and the study by Zhen and colleagues was performed using the Venny online tool^85^.

#### Comparisons to Fontana et al., 2023 characterization of ADP-ribosylation in *Drosophila*

In the *Drosophila* ADPr profiling study^36^, 514 ADPr sites on 296 proteins were identified in *Drosophila* S2R+ cell lines. Flybase IDs of the ADP-ribosylated proteins were matched to orthologous human genes using a list of *Drosophila melanogaster-*Human Orthologs generated from the flybase API on 22.05.2023 and downloaded on 23.07.2023 here: https://ftp.flybase.net/releases/FB2023_03/precomputed_files/orthologs/. 219 matching human orthologs were found, and 77 flybase proteins did not match to any human orthologs. To compare the overlap to the proteins identified in this study, gene names based on de-hyphenated Uniprot IDs were used. Therefore, protein isoforms were not considered.

#### Quantification and statistical analysis of the data

Statistical analysis of ADPr MS data, including principal component analysis (PCA) and volcano plot analysis was performed using Perseus software^84^. Ranked Gene Set Enrichment Analysis (GSEA) on differentially expressed proteome data was performed using fGSEA implemented in R^47^. GSEA on ADPr target proteins was performed using Enrichr implemented in the GSEApy python package^86^. Venn diagram overlaps were calculated using Venny 2.1^85^, and Venn diagrams were generated using python scripts. Cytotoxicity curves for Olaparib, Rucaparib, and Talazoparib were calculated and average CC_50_ values were interpolated in Graphpad Prism using the [Inhibitor] vs. normalized response -- Variable slope nonlinear fit formula. Bar plots and pie charts were generated with Graphpad Prism. UpSet plots were generated with the ComplexHeatmap package in R. Box plots, histograms, and heatmaps were generated with custom R scripts. STRING networks were generated using the Cytoscape app (version 3.10.1)^52^. To construct full STRING networks, dehyphenated Uniprot IDs of the ADPr proteins were used, and the confidence score cutoff was set to 0.7. Functional enrichment was performed in STRING. Filtering was done to keep Reactome pathways and STRING clusters, and redundant terms were removed with a redundancy cutoff of 0.8. Sequence motif logos were generated using iceLogo the web app (v2)^87^. For analysis of the sequence context surrounding serine-ADPr sites, sequence windows of length 15 amino acids N-terminal and 15 amino acids C-terminal to the 1515 identified serine-ADPr sites were used as input, with percentage difference as the scoring system and a p-value cutoff of 0.05. The human Swiss-Prot proteome was used as a reference set.

## Acknowledgements

We thank the lab of Michael O. Hottiger for the expression and purification of recombinant human PARG (University of Zurich), and we thank members of the Novo Nordisk Foundation Center for Protein Research (NNF-CPR) Mass Spectrometry Platform, as well as Ivo Hendriks and Patrick Ruether for instrument support and technical assistance. We thank the lab of Jesper Olsen (NNF-CPR) for providing frozen vials of the breast cancer cell lines used in this study. We thank members of the M.L.N lab for their support. This study was supported by NNF-CPR, the Novo Nordisk Foundation (grant agreement numbers NNF14CC0001 and NNF13OC0006477), the Danish Council of Independent Research (2032-00311B), and the Danish Cancer Institute (R325-A18824,) to M.L.N. H.A.A is supported by the Novo Nordisk Foundation Copenhagen Bioscience PhD program (grant agreement number NNF19SA0035440). Z.S. is supported by the Breast Cancer Research Foundation (BCRF-22-159), Kræftens Bekæmpelse (R325-A18809, R281-16566, and R342-A19788), Det Frie Forskningsræd Sundhed og Sygdom (2034-00205B).

## Author contributions

M.L.N., H.A.A., and S.C.B-L. designed the research question. H.A.A. designed and performed wet lab experiments. A.G.P. and Z.S. performed genomic scar score analyses. H.A.A., M.M., and M.L-P. analyzed the data. H.A.A wrote the original draft of the manuscript. H.A.A, M.M., A.G.P., S.C.B-L, Z.S. and M.L.N. reviewed and edited the manuscript.

---

**Supp. Fig. 1.**
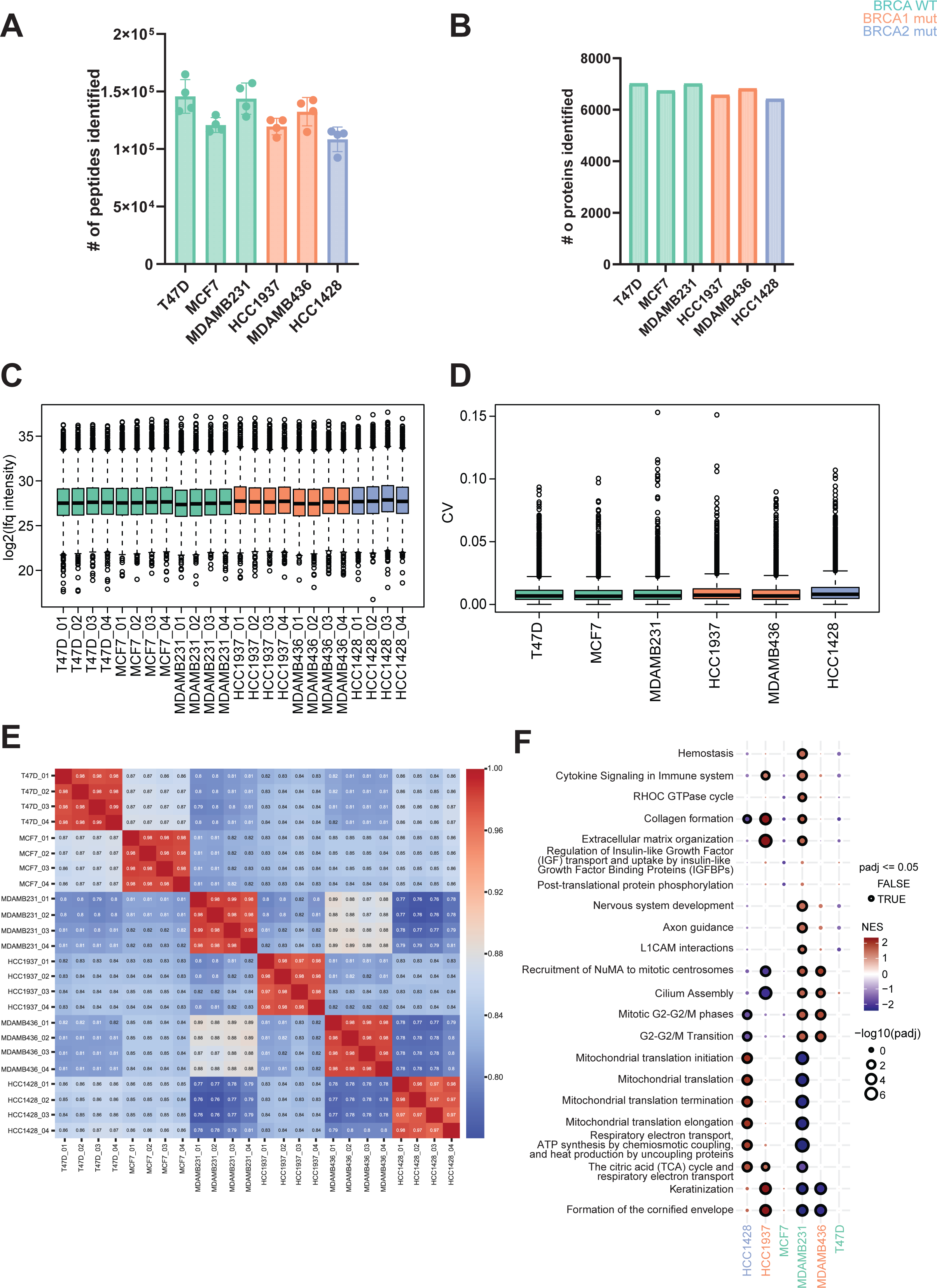

**Supp. Fig. 2.**
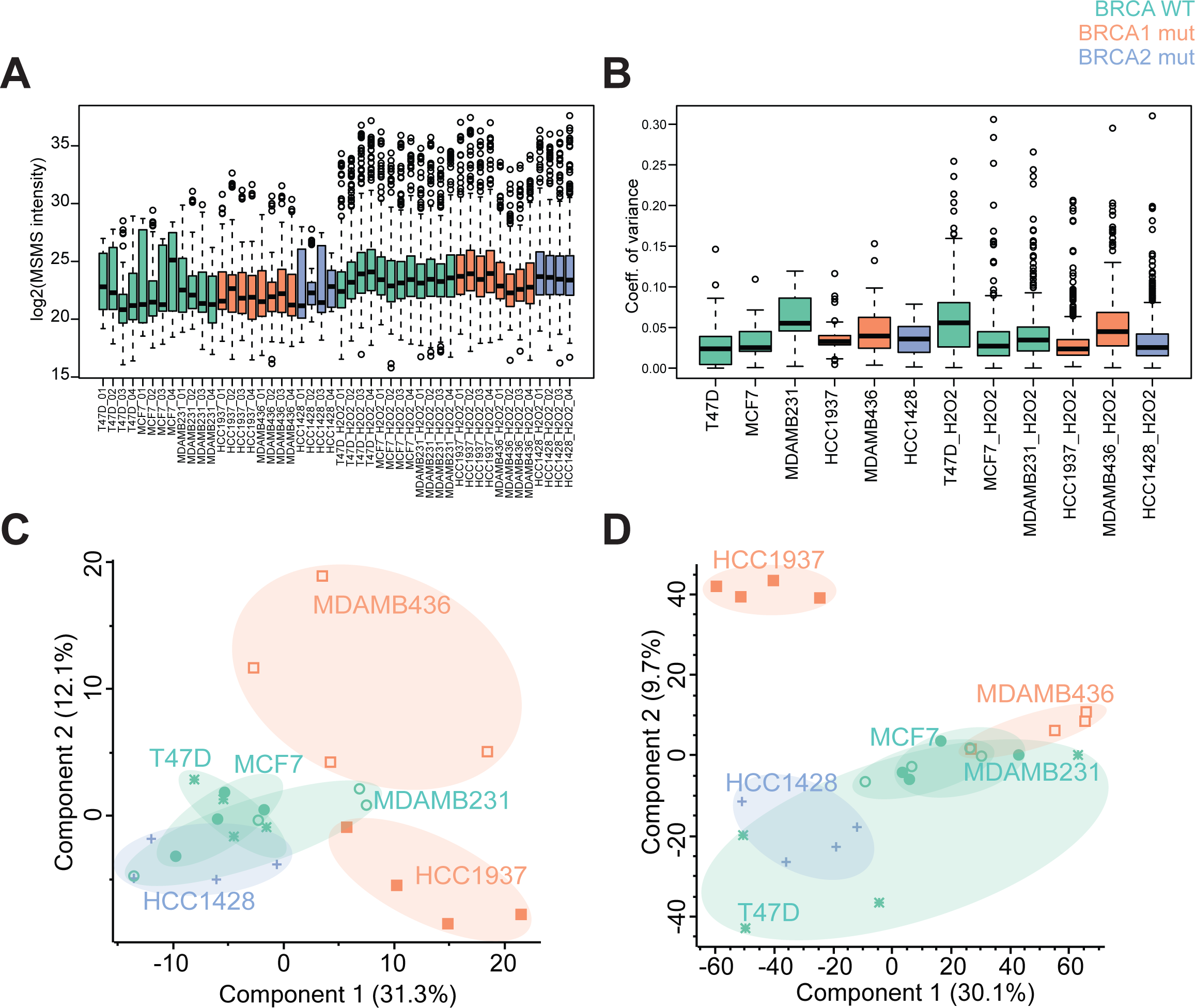

**Supp. Fig. 3.**
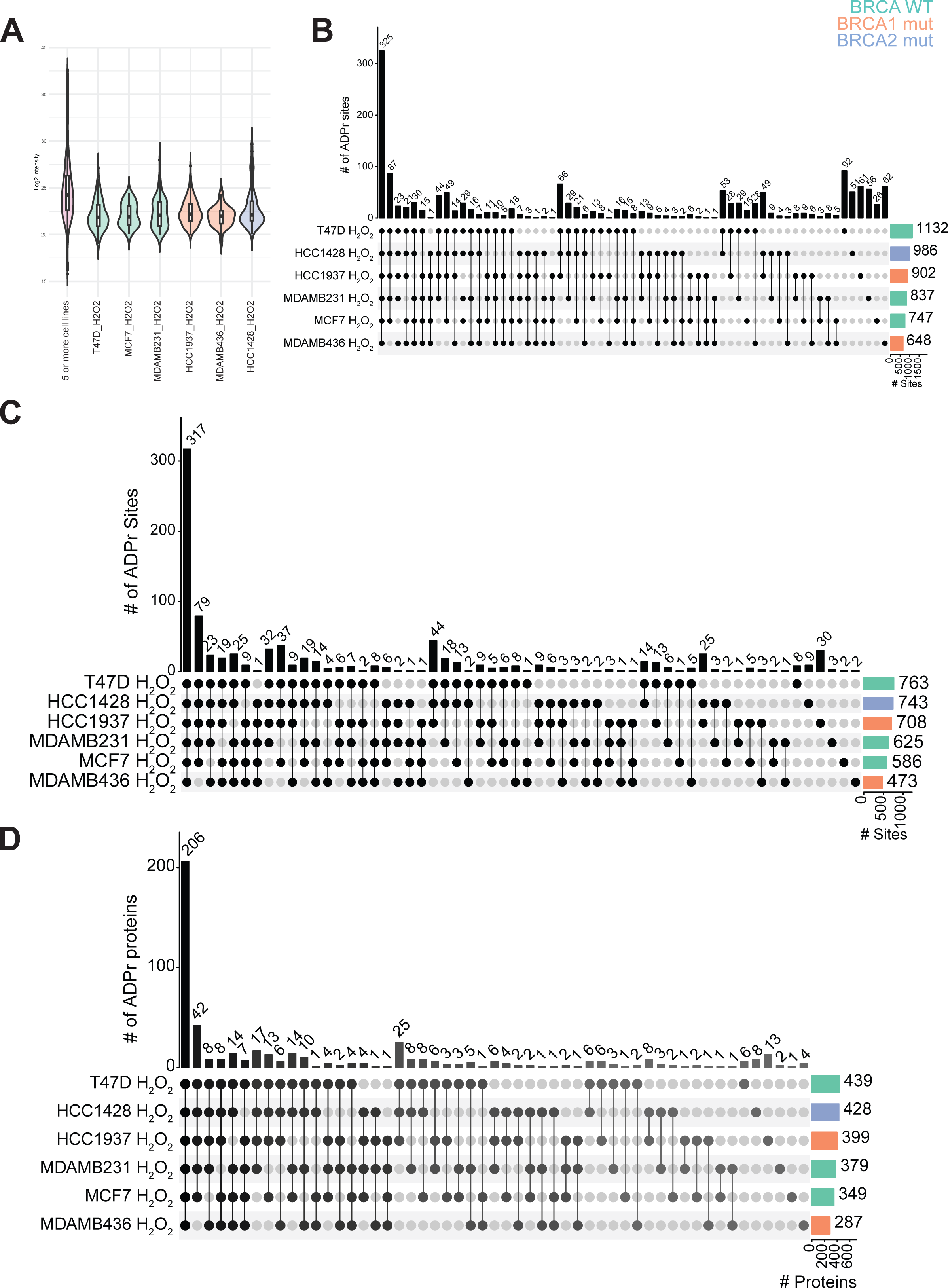

**Supp. Fig. 4.**
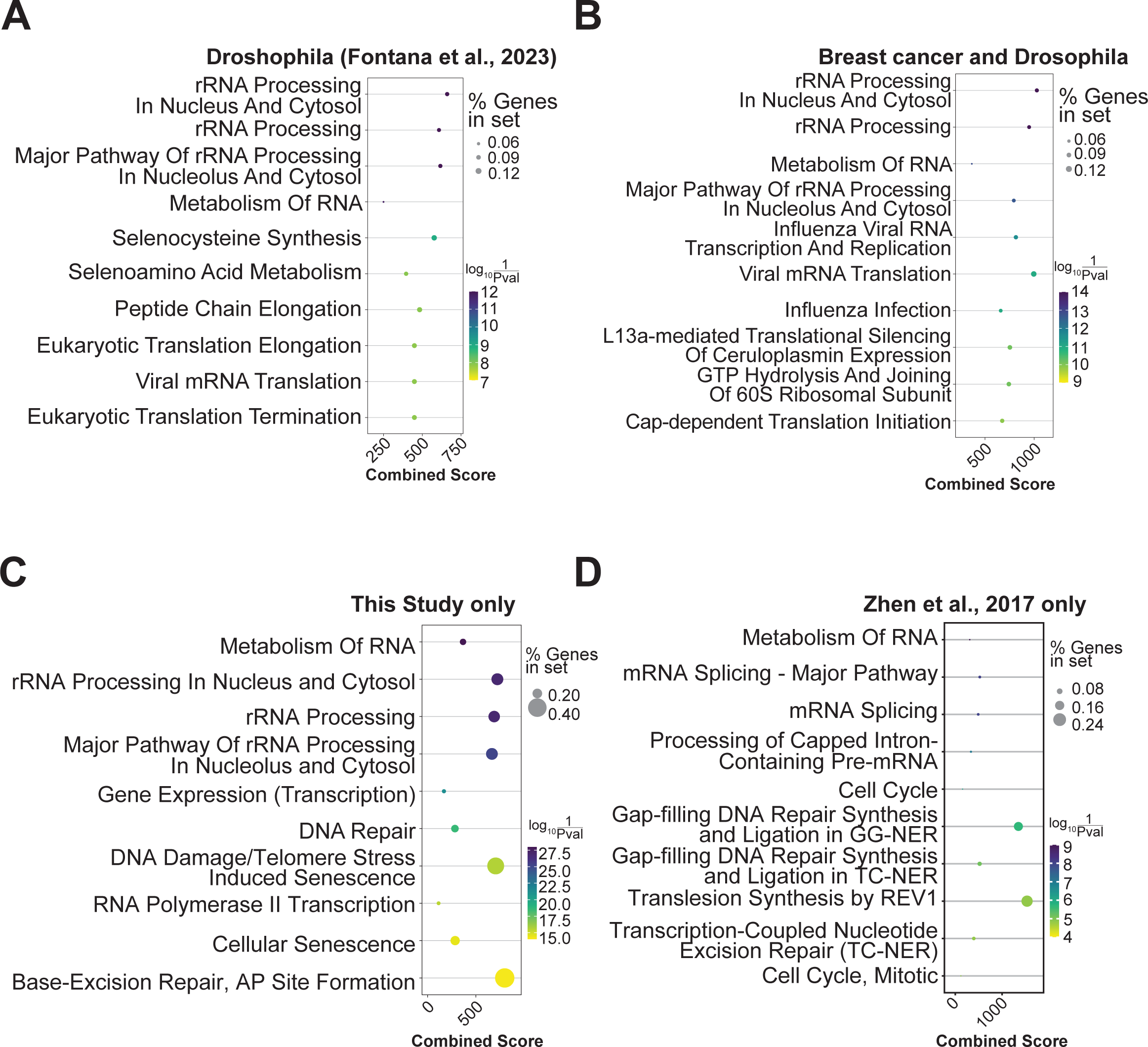

